# DNA-loop extruding condensin complexes can traverse one another

**DOI:** 10.1101/682864

**Authors:** Eugene Kim, Jacob Kerssemakers, Indra A. Shaltiel, Christian H. Haering, Cees Dekker

## Abstract

Condensin, a key member of the Structure Maintenance of Chromosome (SMC) protein complexes, has recently been shown to be a motor that extrudes loops of DNA^1^. It remains unclear, however, how condensin complexes work together to collectively package DNA into the chromosomal architecture. Here, we use time-lapse single-molecule visualization to study mutual interactions between two DNA-loop-extruding yeast condensins. We find that these one-side-pulling motor proteins are able to dynamically change each other’s DNA loop sizes, even when located large distances apart. When coming into close proximity upon forming a loop within a loop, condensin complexes are, surprisingly, able to traverse each other and form a new type of loop structure, which we term Z loop – three double-stranded DNA helices aligned in parallel with one condensin at each edge. These Z-loops can fill gaps left by single loops and can form symmetric dimer motors that reel in DNA from both sides. These new findings indicate that condensin may achieve chromosomal compaction using a variety of looping structures.

---

Control of the spatial organization of chromosomes is critical to life at the cellular level. Structural Maintenance of Chromosomes (SMC) complexes such as condensin, cohesin, and the Smc5/6 complex are key players for DNA organization in all organisms^2–5^. Whereas cohesin is important for gene regulation, DNA damage repair, and holding together sister chromatids^6^, condensin is responsible for the formation of compact mitotic chromosomes and the resolution of sister chromatids in preparation for cell division^7^. An increasing amount of evidence suggests that the underlying principle of DNA organization by SMC complexes is to actively create and enlarge loops of double-stranded DNA, a process named loop extrusion^6^. Polymer simulations^8, 9^ and chromosome conformation capture (Hi-C) data on topologically associating domains^10–13^ suggested the formation of such DNA loops, while recent *in vitro* single-molecule studies provided clear experimental evidence of condensin’s DNA translocase activity^14^ and its ability to extrude loops of DNA^1^.

It remains to be seen how DNA loop extrusion by individual condensins relates to the physiology of highly packed DNA structures. Current modelling has so far assumed that translocating SMC complexes block when they collide, resulting in a string of loops that are clamped together at their stems by adjacent condensins^10, 13^. Experimental evidences for both condensin^15^ and cohesin^16–19^. suggested mutual interactions and a close spacing of SMC proteins^20^. Recent polymer simulations^21^, however, showed that this assumption fails to explain the high degree of compaction observed in mitotic chromosomes^22^ if considering asymmetric extrusion of loops by condensin, the property found in *in vitro* experiments^1^. These findings raise a range of intriguing questions on how these loop extruders orchestrate their actions to organize DNA into complex chromosomes: Do they indeed function independently as individual complexes, or do they exhibit forms of cooperative behavior? What happens when one DNA-loop-extruding SMC complex encounters another one? How do such collisions impact DNA organization? To resolve these questions, we here study the cooperative action of condensin complexes by time-lapse single-molecule visualization. The data reveal a set of distinct interactions between DNA loop-extruding condensin complexes, including the re-shuffling of individual loop sizes and the striking ability of condensin complexes to traverse one another to form a dimeric motor that reels in DNA from both sides and create a novel type of condensed DNA.

To study the interaction between multiple condensin-mediated DNA loops, we imaged the extrusion of DNA loops by budding yeast condensin complexes on 48.5-kilobasepair (kbp) λ-DNA substrates that were tethered at both ends to a passivated surface with sufficient slack to allow loop formation. In these experiments, we labeled the DNA with the fluorescent dye Sytox orange^1^ (SxO) (Fig. 1a). Upon addition of condensin in the presence of ATP, we observed DNA loops as bright fluorescent spots (Fig. 1b). Stretching of these spots within the imaging plane by applying buffer flow perpendicular to the attached DNA confirmed that these spots corresponded to DNA loops (Fig. 1b). While our previous study^1^ focused on the properties of single DNA loops at a concentration of condensin of 1 nM, we here explored slightly higher protein concentrations (2–10 nM), where most DNA molecules displayed multiple loops. Notably, such concentrations, at which we observe a few condensins binding per DNA molecule, approach the in vivo situation where a rough estimate indicates 1 condensin per ∼10 kbp of DNA if all condensins were to bind to the nuclear DNA^23, 24^.

**Figure 1.**
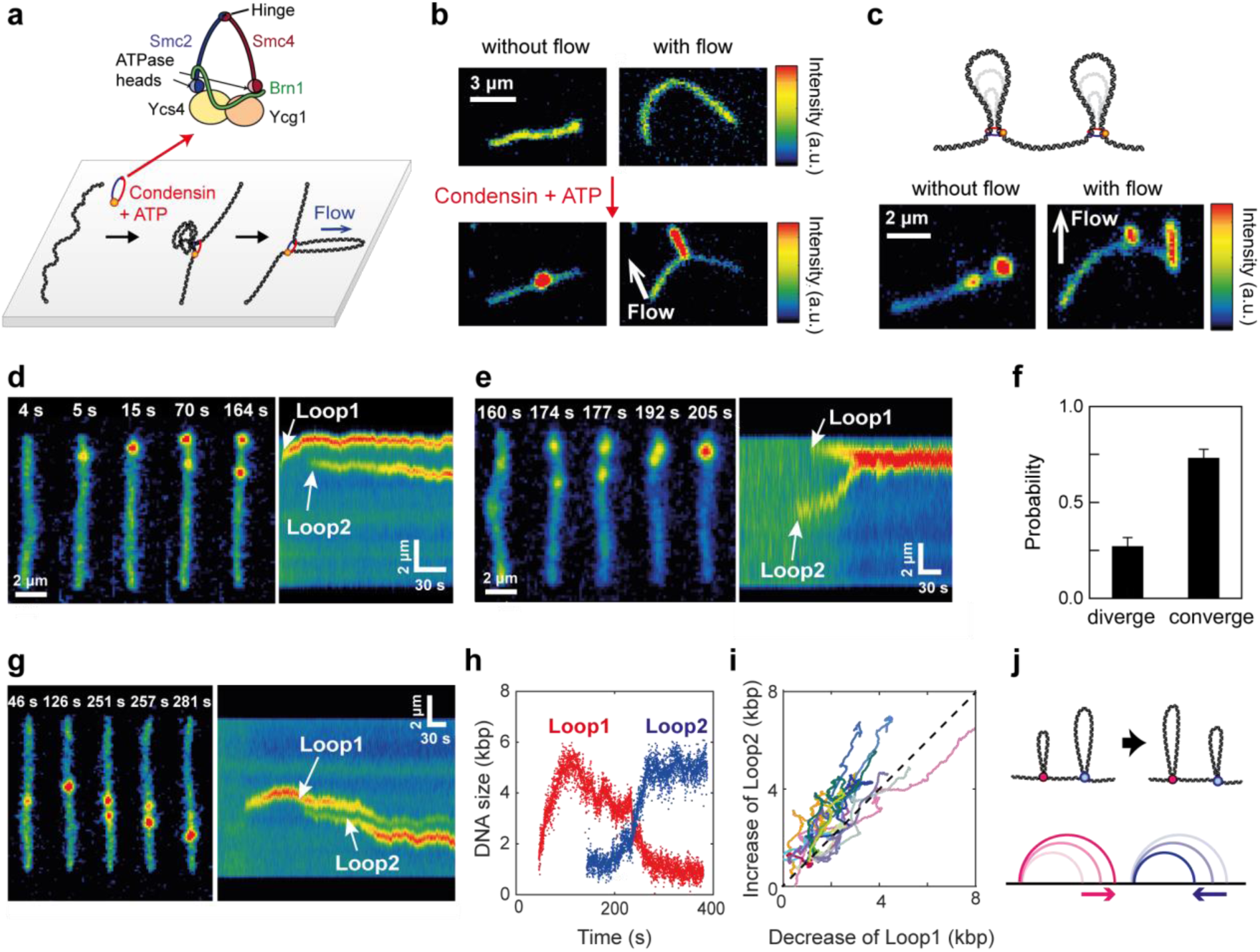
Interactions between multiple condensin-mediated DNA loops. **a**, Cartoon of the *S. cerevisiae* condensin complex which features a large (∼50 nm) ring structure (top), and schematic (bottom) and snapshots **b**, showing single-molecule visualizations of DNA loop extrusion on double-tethered SxO-stained DNA. Arrow in (b) indicates direction of buffer flow. **c**, Schematic and snapshots of two separate loops along a DNA molecule. **d**, **e**, Snapshots (left) and fluorescence-intensity kymographs (right) of two DNA loops that diverge (d) or converge (e). **f**, Probability that two loops are diverging or converging. **g**, Snapshots and kymograph showing the initial formation and shrinkage of a first loop (Loop 1) upon initiation of a second loop (Loop 2). **h**, Corresponding DNA size changes of the two loops in panel (g) versus time. **i**, Simultaneous change of DNA loop size for Loop2 versus for Loop1 (N=16). Dashed line has slope 1, indicating that Loop2 grows at the expense of a shrinkage of Loop1. **j**, Schematic diagram depicting DNA size exchange between two DNA loops in real space (left) and in one-dimensional genomic space (right)^21, 26^.

We first consider the case where two condensin complexes bound at different positions along the same DNA molecule and independently created and expanded loops. In this case, we observed two local maxima along the otherwise homogenous fluorescence intensity of the DNA molecule, which could be stretched into loops under buffer flow (Fig. 1c). Since yeast condensin extrudes DNA loops asymmetrically^1^, where the side from which DNA is reeled into the loop is presumably determined by the orientation of the Ycg1/Brn1 DNA-anchor site to the DNA^1, 25^, two individual DNA loops either moved into opposite directions (Fig. 1d) or towards each other (Fig. 1e). We observed a 27/73% distribution of mutually diverging or converging loops (Fig. 1f, N = 60), in a good agreement with the expected 25/75% distribution for a random orientation of the condensins, given that one out of four possible orientations of two condensin complexes (anchor sites facing towards each other) gives rise to diverging loops, while the other three orientations lead to converging loops.

We find that loops can influence each other even if separated far apart. In most cases, loops on a DNA molecule started to form at different time points (e.g. Fig. 1g), confirming that loops initiate independently of other loops on the same DNA. Remarkably, upon initiation of a second loop, the pre-existing loop frequently began to shrink (70% of cases; N = 40) (Fig. 1g, Supplementary Video 1). The changes in DNA length of the two loops, estimated from fluorescence intensities, showed a clear anticorrelation (Figs. 1h and 1i and Extended Data Fig. 1, N=17). In other words, the new DNA loop extruded by the second condensin grew at the expense of the original one, which concomitantly shrank in size over time. This indicates that DNA in a loop can slip back through the condensin motor, likely because of an increase in DNA tension that occurs as the second condensin docks and starts reeling in DNA. The finding that docking and loop extrusion of a remotely located condensin on the same DNA substrate can induce shrinkage of an already extruded loop is unexpected and implicates that it is possible to redistribute individual loop sizes (Fig. 1j).

Surprisingly, these separate individual loops were *not* the majority class of DNA structures in these experiments. Instead of independent parallel DNA loops, we predominantly observed a higher-order DNA structure that appeared as an elongated line of high fluorescence intensity stretching along the length of DNA (Fig. 2a). Imaging under a sideways flow revealed that the observed structure consisted of three dsDNA stretches that were connected in parallel (Fig. 2a, Extended Data Fig. 2 and Supplementary Video 2). We name this structure a *Z loop*, since the shape of this loop resembles the letter Z. The probability to observe Z loops increased with the protein concentration and became the majority pattern for condensin concentrations higher than 6 nM (Fig. 2b).

**Figure 2.**
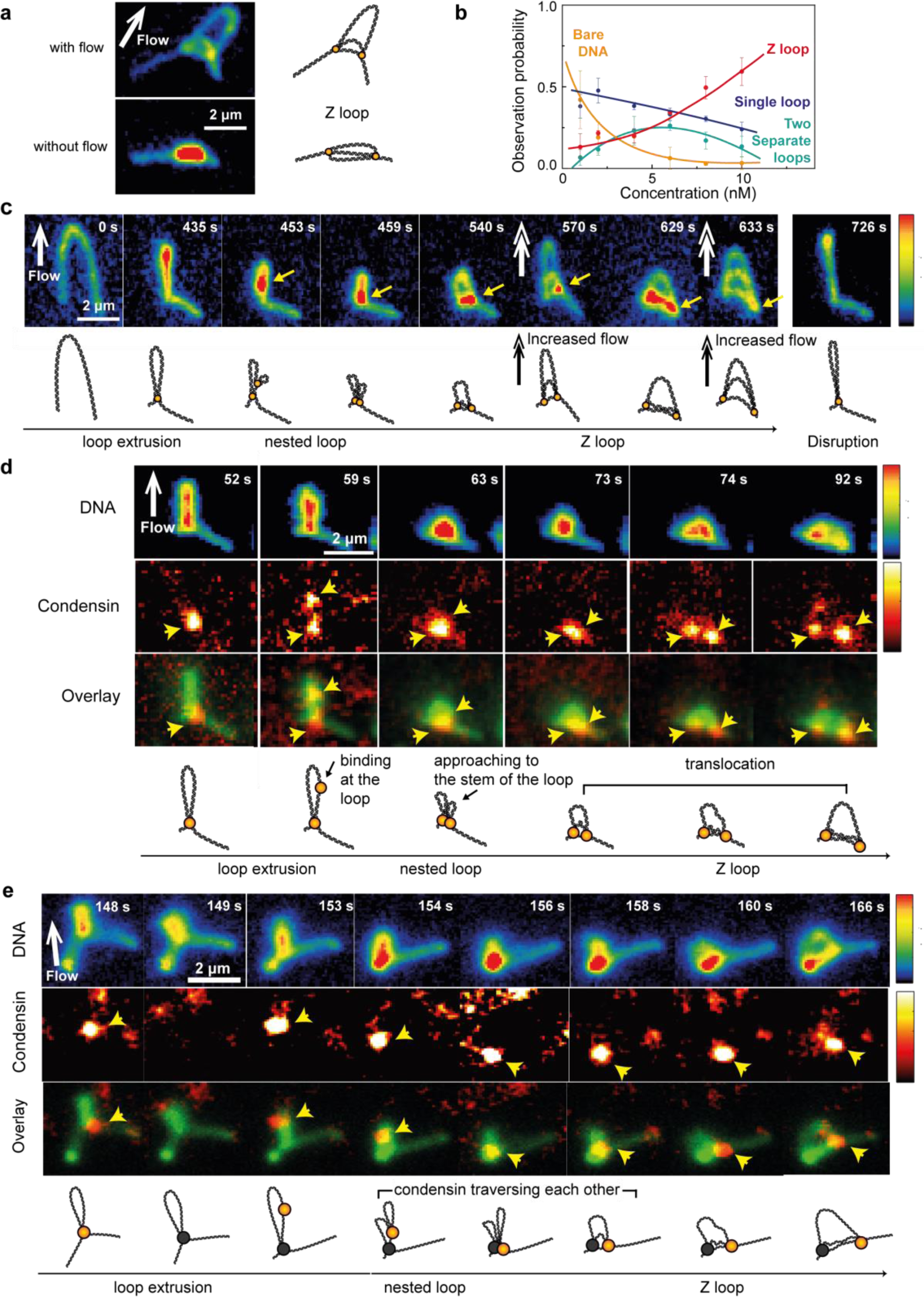
Condensins can traverse one another and form a Z loop on DNA. **a**, Schematic and snapshot observations of a Z loop, a linearly stretched DNA structure along the DNA molecule that consists of three dsDNA molecules in parallel, as revealed by application of an inplane buffer flow (white arrow). **b**, Probability of observing DNA molecules exhibiting no loop (black), a single loop (blue), two separate loops (turquoise), and a Z loop (red), as a function of condensin concentration. Error bars represent SD from four different data sets (N_tot_=476 molecules). Lines serve as guides to the eye. **c**, Series of snapshots showing DNA intermediates in Z-loop formation. Initially, a single loop is extruded which enlarges until it reaches the tethered end of DNA at t = 435 s. Subsequently, a locally compacted region appears and approaches to the stem of the initial loop (∼459 s), followed by a gradual elongation into a Z loop (633 s). Occasionally the buffer flow rate was increased (570s, 633s), which revealed that the Z loop was indeed composed of three parallel DNA segments (540s, 629s). This Z loop finally disrupts into a single loop (726 s). Yellow arrows denote the moving DNA parts. Schematic diagrams underneath the images provide visual guidance. **d**, Snapshots of SxO-stained DNA (top), ATTO647N-labeled condensin (middle), and their overlay (bottom) showing the locations of two condensins and the simultaneous transitions of DNA conformations during Z loop formation. A second condensin binds at the location near the loop that is created by the first condensin (59 s), and approaches to the first one while compacting the single loop further (63 s), whereupon one of the two condensins moves away from the other one as Z loop extends (92 s). Schematic diagrams underneath the images provide visual guidance. Arrows denote the location of the two condensins. **e**, Snapshots of DNA (top), condensin (middle), and their overlay (bottom) tracing the locations of the second condensin during Z loop formation. The fluorescence signal from the first condensin that extruded the single DNA loop (148 s) was photobleached (149 s) before the binding of the second condensin at the DNA-loop region (153 s). This condensin again approaches to the loop stem (156 s), followed by the linear translocation along the DNA outside of the initial loop (∼166 s), where it shortly paused, and subsequently linearly translocated along the DNA outside of the initial loop (158-166s). Schematic diagrams underneath the images provide visual guidance. Yellow arrows denote the location of the second condensin.

Real-time imaging of the flow-stretched DNA revealed the characteristic formation of a Z loop (see Fig. 2c and Supplementary Video 3 for a typical example): After a single loop extruded (t=435s) and stalled^1^, a locally compacted region – presumably a small loop formed by an additional condensin – appeared within the initial loop (453s) and approached to the stem of the single loop (459s). This ‘nested loop’ of two smaller parallel loops did not, as could be expected, stop at this point, but instead began to extend towards the DNA outside of the initial loop (540s) and continued to stretch until it either hit the tethered end of DNA (629s) or until one or more of the three dsDNA stretches were fully extended (see Extended Data Fig. 3 for additional examples). To trace the position of the condensin complex during the transition from single loop to nested loop to Z loop, we co-imaged DNA and condensin labelled with a single fluorophore (ATTO647N) (Fig. 2d and Supplementary Video 4). This revealed that after some time Δt_1_ after the initiation of the first loop, an additional condensin complex bound to a position within the initial DNA loop (59s) and subsequently approached the stem of the loop (63s), where the first condensin was located. Simultaneously, the conformation of DNA changed from a single loop (59s) into a more compacted form (63s), indicating the formation of a nested loop extruded by the additional condensin. After a brief waiting time Δt_2_, one of these condensins then moved away from the stem of the loop and linearly translocated along the DNA outside of the loop. In doing so, the nested loop changed its conformation to a Z loop, with the two condensins localized at both edges of the Z loop (92s) (see Extended Data Fig. 4 for additional examples). To identify which of the two condensins co-localized at the ‘leading edge’ of the Z loop, we examined events where the first condensin had bleached before the binding of the second condensin (Fig. 2e, Extended Data Fig. 5, Supplementary Videos 5 and 6). These experiments showed that it was the second condensin that, strikingly, traversed the first condensin at the base of the first DNA loop.

**Figure 3.**
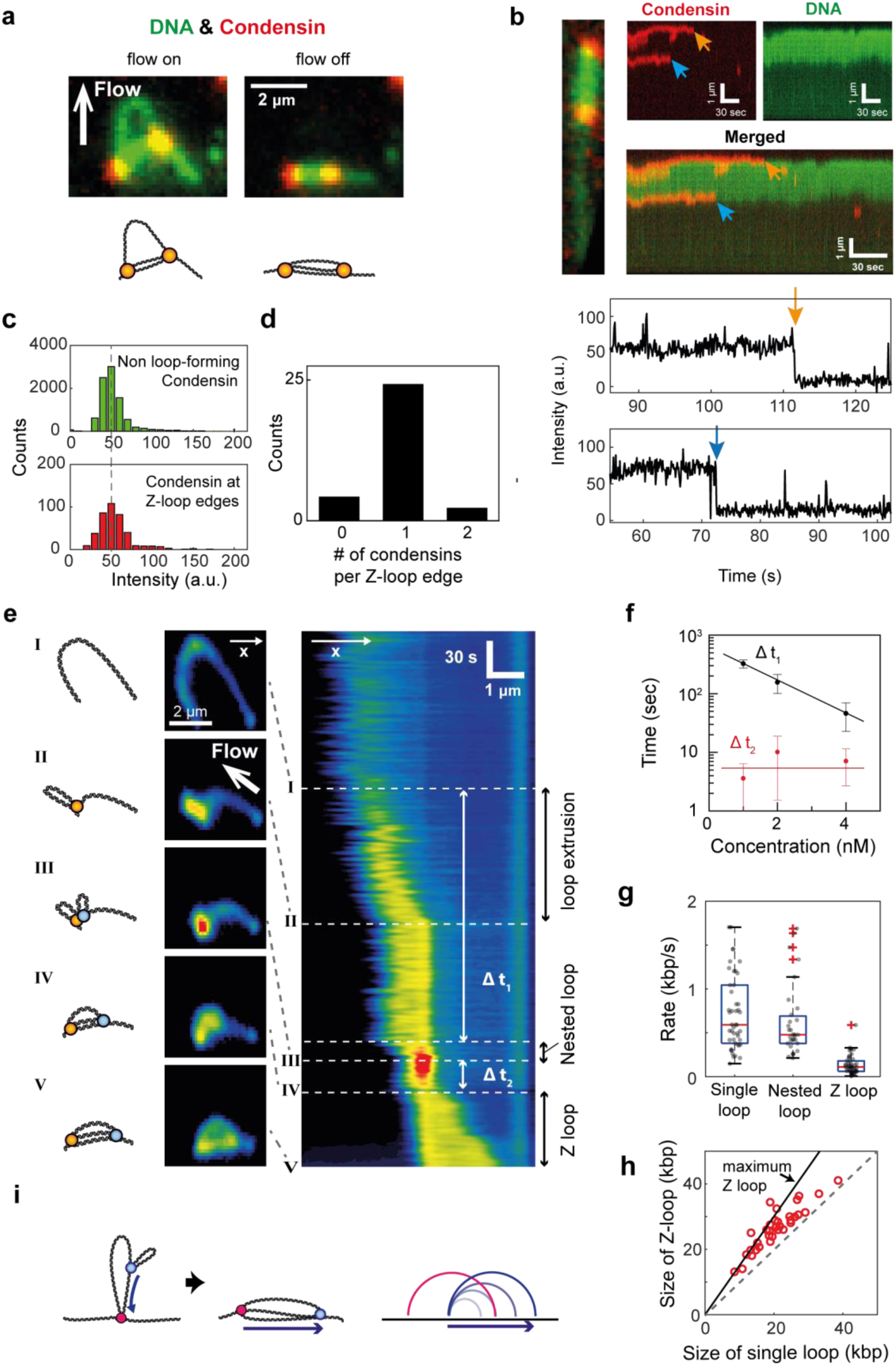
Quantification of Z loop formation kinetics. **a**, Overlayed images of SxO-stained DNA and ATTO647N-labeled condensin showing a Z loop in the presence (left) and the absence (right) of buffer flow. For the quantification of number of condensins, Z loop formation was monitored in the absence of flow. **b**, Snapshot and kymographs (top) of SxO-stained DNA (green), ATTO647N-condensin (red), and their merge, and corresponding ATTO647N fluorescence time traces (bottom) showing single-step photobleaching events from the condensins bound at each edge of the Z loop. **c**, Fluorescence intensity distributions for condensins that are temporarily bound on DNA but did not lead to loop extrusion (top), and for those that are located at Z-loop edges (bottom). **d**, Number of condensin complexes at each edge, estimated from the two histograms of (C). **e**, Fluorescence intensity kymograph along the axis connecting both ends of the DNA tether showing a transition of DNA conformation from a bare DNA, to single loop, to nested loop, to a gradual extension of a Z loop, and the related time lags. **f**, Waiting time between starting the single loop and starting the nested loop (Δt_1_), and between the start of the nested loop and the Z loop (Δt_2_), plotted against the protein concentrations. Lines serve as guides to the eye. **g**, Rate of single loop, nested loop, and Z loop formation, estimated from fluorescence kymographs such as (e) (see Methods). **h**, Final Z-loop size plotted verses the single-loop size before Z-loop formation. A linear relation is observed, with 30% of molecules showing the maximum possible sizes of Z loop which is 1.5 times that of the initial single loop (N=32). **i**, Schematic diagram depicting Z-loop formation depicted in real space (left) and one-dimensional genomic space (right).

**Figure 4.**
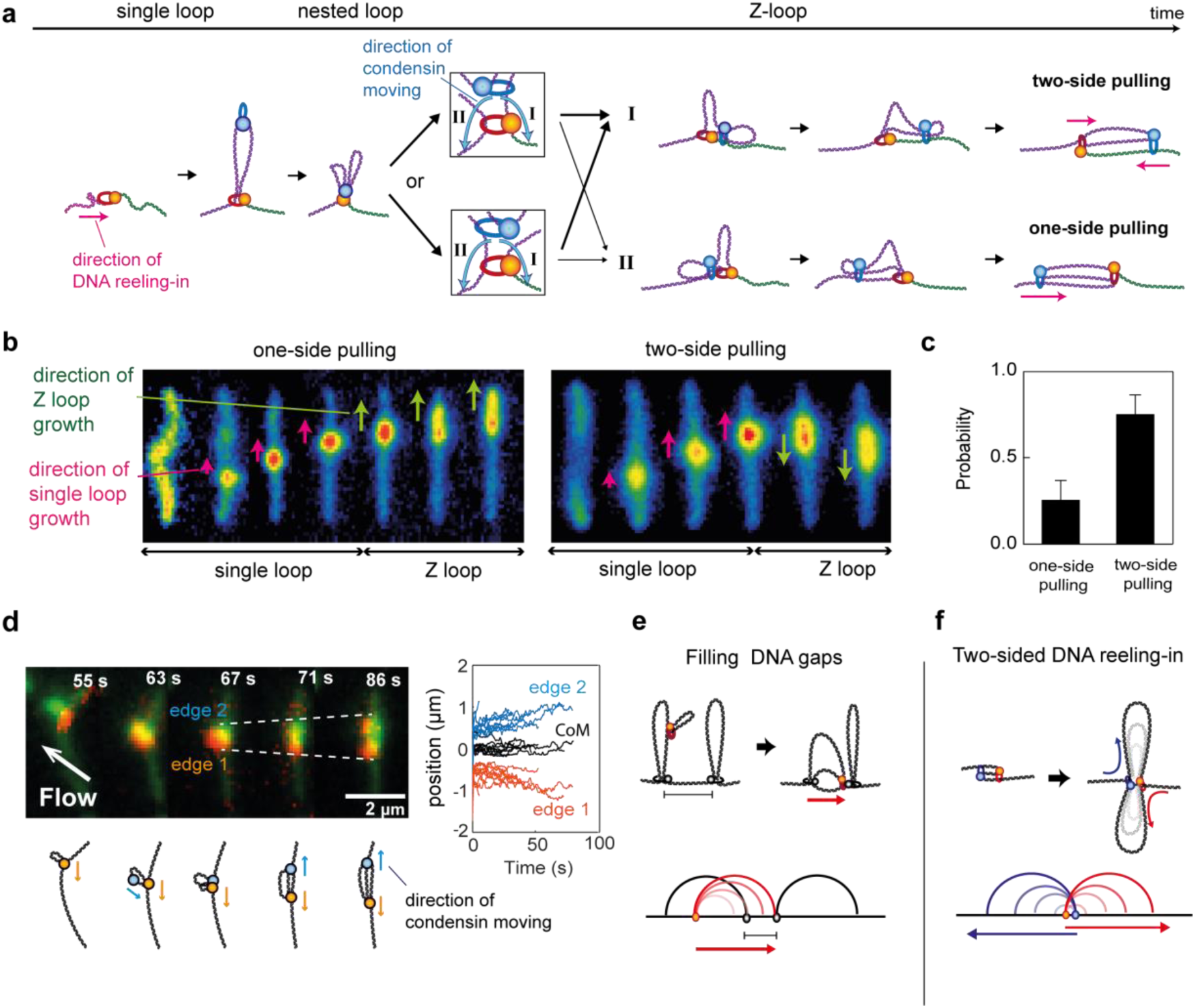
Possible impact of Z loops on chromosomal compaction. **a**, Model of DNA Z-loop formation by two condensins. Depending on the orientations of the two condensins (see zooms), the formed Z loop can reel in DNA either from both sides of DNA (two-side pulling) or from one side of DNA (one-side pulling). **b**, Series of snapshots showing two DNA molecules where the initial single loop and the subsequent Z loop grow from the same side of DNA (left) or from opposite sides (right). **c**, Probability that a Z loop pulls from one side or from two sides. **d**, Snapshots (top left) and schematics (bottom left) of overlay of SxO-stained DNA and ATTO647N-labeled condensin. For this molecule, binding of the second condensin occurred before the first condensin fully extruded the single loop, thus allowing for the first condensin to continue to reel in DNA during Z loop extension. This results in a symmetric divergence of the two condensins. Simultaneous change of positions of two Z loop edges (blue and orange) and of the center of mass (black) (N=11; right). **e**, **f**, Schematic diagrams depicting possible implications of Z loops for chromosomal compaction depicted in real space (top) and one-dimensional genomic space (bottom).

We then quantified the data of these experiments. Photobleaching (Figs. 3a and 3b, Extended Data Fig. 6, Supplementary Video 7) and intensity distributions (Figs. 3c and 3d) confirmed that indeed two single condensin complexes, i.e. not dimers or multimers, were involved in the Z-loop formation (Fig. 3d). Fluorescence intensity kymographs (Fig. 3e and Extended Data Fig. 7) revealed the clear transition of the DNA conformations: from bare DNA, to extrusion of a single loop, to formation of a nested loop, to linear extension into a Z loop. We observed that the initial lag time Δt_1_, the interval between the start of the initial loop and the start of the nested loop, decreased with the protein concentration (Fig. 3f, N=31), as expected, since it essentially equals the time lag between binding events of the first and second condensin. The second lag time Δt_2_, the interval between the end of nested loop formation and the start of Z loop formation, however, was independent of protein concentration (7 ± 6 s, mean ± SD, N=31). This quantifies the time that two condensins spent in close proximity to each other before the second condensin traversed the first one. The loop-formation rate was similar for single loops and nested loops (Fig. 3g and Extended Data Fig. 8, see Methods): 0.7 ± 0.4 kbp/s (mean ± SD, N=46) and 0.6 ± 0.4 kbp/s (mean ± SD, N=31), respectively. This is consistent with the notion that the observed compaction of the single loop is induced by a second condensin creating, at the same speed, a loop within the initial loop. The observed rate of Z-loop formation, however, was much lower, only 0.1 ± 0.1 kbp/s (mean ± SD, N=47), which likely is due to the different mechanism of Z-loop formation, where condensin translocates along DNA while dragging the nested loop alongside. Once formed, Z loops were found to be very stable, even more so than single loops (Extended Data Fig. 8). The length of Z loops was linearly proportional to the length of the initial single loop (Fig. 3h, N=32), with a good fraction (∼30 %) exhibiting close to the maximum possible length (∼1.5 *l*), that is the length of the initial loop (1.0 *l*) plus the maximum possible distance between two condensin anchors, i.e. 0.5 *l* for the case of the second condensin binding at the tip of the initial loop (Extended Data Fig. 9). This implies that a single Z loop does not linearly compact DNA more when compared to two individual loops (see Extended Data Fig. 9), but rather changes its topology into a more condensed structure with connected DNA-loop and non-loop-DNA stretches (Fig. 3i).

Our data reveal the characteristic pathway of two condensins that traverse each other and, as a result, form a Z loop (Fig. 4a). Upon forming a loop within a loop, the second condensin comes into close proximity of the first condensin, shortly pauses (Δt_2_ ∼ 7s), and then reaches out to the DNA outside of the first loop. Here, regardless of the two possible relative orientations of two condensins (zoomed images in Fig. 4a), the second condensin can in principle reach out to the DNA next to either the anchor or the opposite site of the first condensin (route I or II, respectively). After the direction is chosen, the second condensin traverses the first condensin and translocates along the DNA, dragging along its own loop, and forming a three-stranded Z loop. Interestingly, as can be seen from the right-side drawings in Fig. 4a, these two routes lead to qualitatively different loops, viz., a Z loop that reels in DNA from both sides (top) or one side (bottom). By comparing the relative direction of single-loop growth before stalling and the direction of subsequent Z loop extension (Fig. 4b), we, surprisingly, did not observe a 50/50% distribution of both types as expected for random relative orientations of the two condensins, but rather a 75/25% distribution of two-side-pulling verses one-side-pulling Z loops (Fig. 4c, N=70). The observed strong bias to form two-side-pulling Z loops likely originates from a preference for the second condensin to traverse to DNA beyond the anchor site of the first condensin (route I). In the (rare) cases that a Z loop initiated very early, i.e., for small Δt_1_ and thus binding of the second condensin before the initial single loop was fully extruded, we observed that the Z loop expanded symmetrically to both directions, directly confirming the two-side pulling (Fig. 4d, N=11, Supplementary Videos 8 and 9). This shows that condensin, which individually is a one-side pulling motor, can cooperatively reel in DNA from both sides in the form of two-side pulling Z loops. Once Z loops were fully extended, they occasionally (∼30%) slipped DNA from one or both of the edges, leading to random diffusion along the DNA tether over time (Extended Data Fig. 10)

These discoveries of interactions between DNA-loop extruding condensin complexes have important implications for understanding the fundamental mechanisms of chromosome organization. The finding that multiple DNA loops can change their sizes via slippage adds valuable information to the current picture of loop-extrusion dynamics, which so far only considered the formation, growth, and dissociation of loops. Most importantly, the unanticipated property of condensin complexes that they can traverse one another has direct consequences for modeling of chromosomes^9, 12^. A consensus view has emerged that mitotic chromosomes consist of an array of individual single loops in parallel. Whereas our previous discovery of an asymmetry of DNA loop extrusion posed a problem, since this mechanism would leave gaps in-between loops^21^, the newly discovered Z loops may extend along the unextruded parts of DNA, thereby filling such gaps (Fig. 4e, top, and Supplementary Video 10). Another scenario may be that Z loops are the norm rather than individual parallel loops, which is likely to occur if condensin prefers to extrude loops within loops rather than at non-looped DNA stretches, for example due to internal forces within the DNA induced by the network topology or active processes in vivo. If Z loops initiate rapidly after the nucleation phase of single loops, they result in two condensins that are anchored close to each other, yielding effectively a two-side-pulling condensin dimer structure that reels in DNA in a symmetric fashion (Fig. 4f, bottom). Models of chromosome compaction will need to consider these findings that SMC proteins exhibit the ability to form a rich variety of looping structures.

## Acknowledgments

We thank S. Bravo, J. van der Torre, and E. van der Sluis for technical support, and J. Eeftens, M. Ganji, A. Katan, B. Pradhan, B. Rowland, J.-K. Ryu, and J. van der Torre for discussions.

## Author contributions

E. K. and C. D. designed the single-molecule visualization assay, E. K. performed the imaging experiments, J. K. contributed in image analyses, I. A. S. developed condensin fluorescence labeling strategies and purified protein complexes, C. D. and C. H. H. supervised the work, all authors wrote the manuscript.

## Data and materials availability

Original imaging data and protein expression constructs are available upon request.

## Methods

### Condensin holocomplex purification

We used our previously published expression and purification procotols^1^ to prepare the pentameric *S. cerevisiae* condensin complex.

### Fluorescent labeling of purified condensin complexes

The purified condensin complexes were fluorescently labeled as described previously^1^. Briefly, a 10 % excess of ATTO647N-maleimide (ATTO-TEC) was coupled to Coenzyme A (Sigma) in deoxygenated 100 mM sodium phosphate buffer at pH 7.00 for one hour at room temperature. 10 % equivalent of tris(2-carboxyethyl)phosphine was included halfway through the reaction and coupling was terminated with an excess of dithiothreitol. The reaction mixture was used for enzymatic covalent coupling to ybbR acceptor peptide sequences within the kleisin subunit in condensin holocomplexes (Brn1[13-24 ybbR, 3xTEV141]-His_12_-HA_3_; C5066), using a 5-fold excess of fluorophore to protein and ∼1 μM Sfp synthase (NEB) for 16 hours at 6 °C in 50 mM TRIS-HCl pH 7.5, 200 mM NaCl, 5% v/v glycerol, 1 mM DTT, 0.01% Tween-20, 0.2 mM PMSF, 1 mM EDTA. Labeled protein was separated from unreacted fluorophore and the Sfp synthase by size-exclusion chromatography on a superose 6 3.2/200 (GE Healthcare) preequilibrated in 50 mM TRIS-HCl pH 7.5, 200 mM NaCl, 5% v/v glycerol, 1 mM MgCl_2_, 1 mM DTT).

### Double-tethered DNA assay for single-molecule imaging

Phage λ-DNA molecules were labelled with biotin at their both ends as described previously^1^. The biotinylated DNA molecules were introduced to the streptavidin-biotin-PEG coated glass surface of a flow cell at constant speed of 5 – 10 μL/min, resulting in attachment of DNA molecules with relative DNA extensions ranging from ∼0.3 to ∼0.6. The surface-attached DNA molecules were stained with 500 nM Sytox Orange (Invitrogen) intercalation dye and imaged in condensin buffer (50 mM TRIS-HCl pH 7.5, 50 mM NaCl, 2.5 mM MgCl_2_, 1 mM DTT, 5% (w/v) D-dextrose, 2 mM Trolox, 40 µg/mL glucose oxidase, 17 µg/mL catalase).

Real-time observation of multiple loop interactions by condensin was carried out by introducing condensin (1-10 nM) and ATP (5 mM) in the above specified condensin buffer. Although Z-loops were more frequently observed at higher concentrations (6-10 nM), most of the presented data were obtained in the concentration range of 2-4 nM. This was done to study single Z loops and minimize measuring on DNA molecules that exhibited both a single loop and Z-loop simultaneously. For dual-color imaging of SxO-stained DNA and ATTO647N-labeled condensin, we also kept to this lower concentration range to minimize the background coming from both freely diffusing labelled condensin as well as from labelled condensins that transiently bound onto DNA without forming DNA loops.

Fluorescence imaging was achieved by using a home-built epi-fluorescence/TIRF microscopy. For imaging of SxO-stained DNA only, a 532-nm laser was used in epi-fluorescence mode. In the case of dual-color imaging, SxO-stained DNA and ATTO647N-labelled condensin were simultaneously imaged by alternating excitation of 532-nm and 640-nm lasers in Highly Inclined and Laminated Optical sheet (HILO) microscopy mode with a TIRF objective (Nikon). All images were acquired with an EMCCD camera (Ixon 897, Andor) with a frame rate of 10 Hz.

### Data analysis

Fluorescence images were analysed using custom-written Matlab (Mathworks) software. Fluorescence intensity kymographs as shown in e.g. Fig. 1d were built from the intensity profiles of DNA molecules per time point as explained in our previous paper^1^. From the kymographs for individual molecules thus obtained, a loop analysis (cf. Figs. 1h, Extended Data Fig. 1) was carried out as follows. The start center position and start time of a loop was indicated through user input. Then, the position of the loop at each time point was found by center-of-mass tracking over a section of the DNA molecule. This procedure was repeated until a user-set end time for this loop was reached. The data was stored as a position-time trace per loop.

The ‘left’ and ‘right’ position of the loops were used to define three intensities:

1. ‘Left’: intensity from the section left of the left loop edge (region I).
2. ‘Middle’, comprising the remainder of the intensity counts: the loop excess counts plus the tether section in between the loop edges (region II).
3. ‘Right’: intensity from the section right of the right loop edge (region III).

These three intensities were then expressed as percentages of the total intensity count, adding up to 100%. Using this intensity information, the sizes of DNA (in kbp) within individual single loops (e.g. Fig. 1h) and the size of DNA within Z loops (e.g. Fig. 3h), were obtained by multiplication of the percentages in the loop region II by 48.5 kbp.

To compare the rates of single loop extrusion, nested loop formation, and Z loop extension we first built the intensity kymographs during Z loop formation under buffer flow application (e.g. Fig. 3e and Extended Data Fig. 8). Note that the change from a single loop to a nested loop can only be seen by flow-induced DNA stretching. From the kymographs, the rate of nested loop formation is determined by the change in the physical length of single loop and that of nested loop, divided by the time duration. Note that unlike for nested loop formation, where there is no internal force within the loop, the single and Z loop formation is influenced by the tension within the DNA tether, complicating a direct comparison of the three rates. Therefore we estimated the force applied to DNA during buffer flow application by measuring the relative extension of single-tethered lambda DNA molecules under flow application, which yielded a value of ∼0.2-0.3 pN. Then, we extracted the rates of single loop and Z loop formation by turning off the buffer flow and monitoring molecules for which the initial applied force value was in a similar range. The time traces of DNA size changes in the loop region during Z loop formation (e.g. Extended Data Fig. 9) among those molecules were then used to determine the rates. For the extraction of the respective rates, a simple linear fit to the linear increase of DNA amount during the first 10 seconds of the initial single loop and the following Z loop growth was used. In this way, the rates of single-loop, nested-loop and Z-formation were obtained under similar conditions of an applied force of 0.2-0.3 pN.

## Extended Data Figures

**Extended Data Figure 1.**
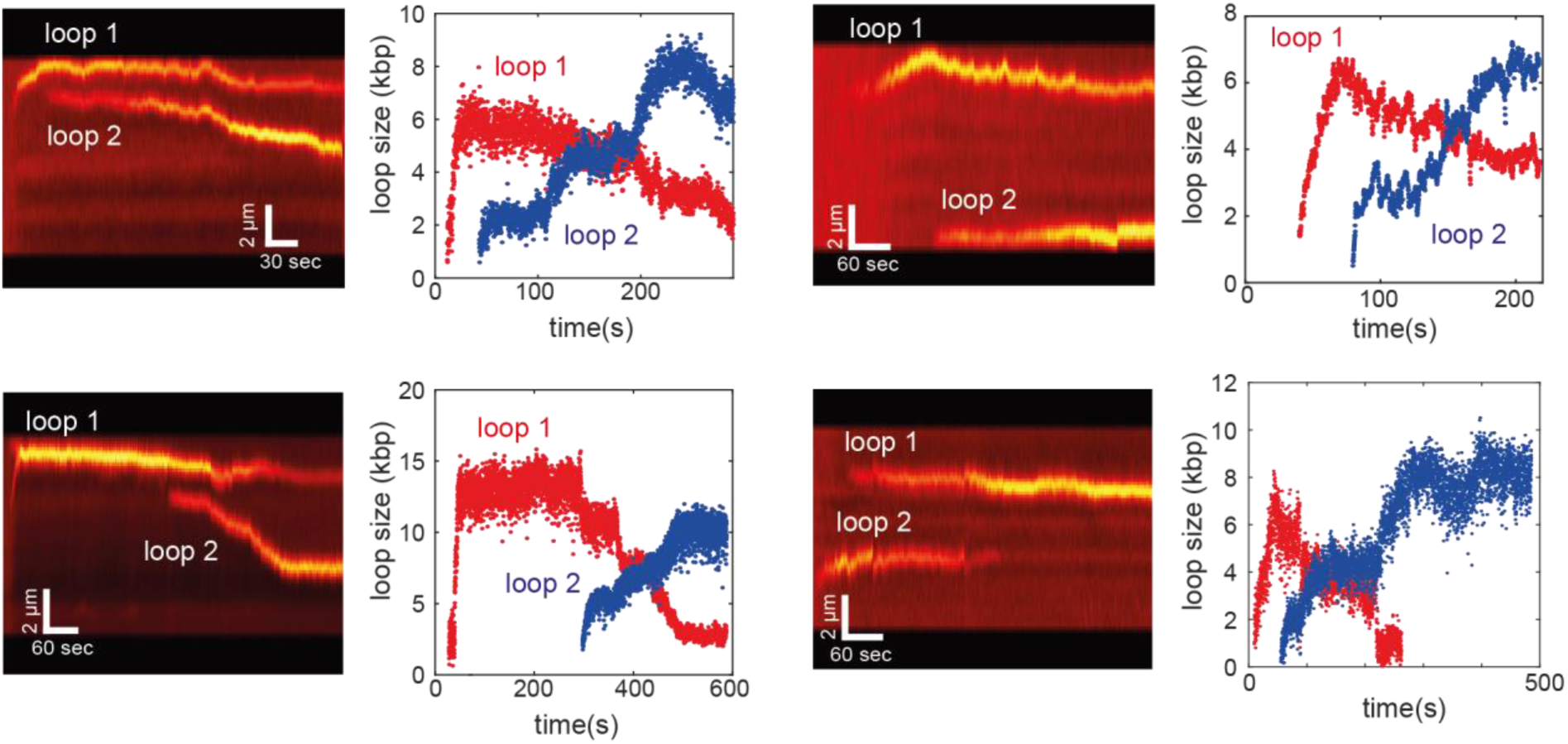
Additional examples of kymographs showing a redistribution of loops sizes. Left panels are kymographs that show a shrinkage of an initially formed loop (Loop 1) upon initiation of a second loop (Loop 2), and right panels are the corresponding DNA size changes between the two loops, calculated from the integrated fluorescence intensities and the known 48.5-kbp length of the λ-DNA.

**Extended Data Figure 2.**
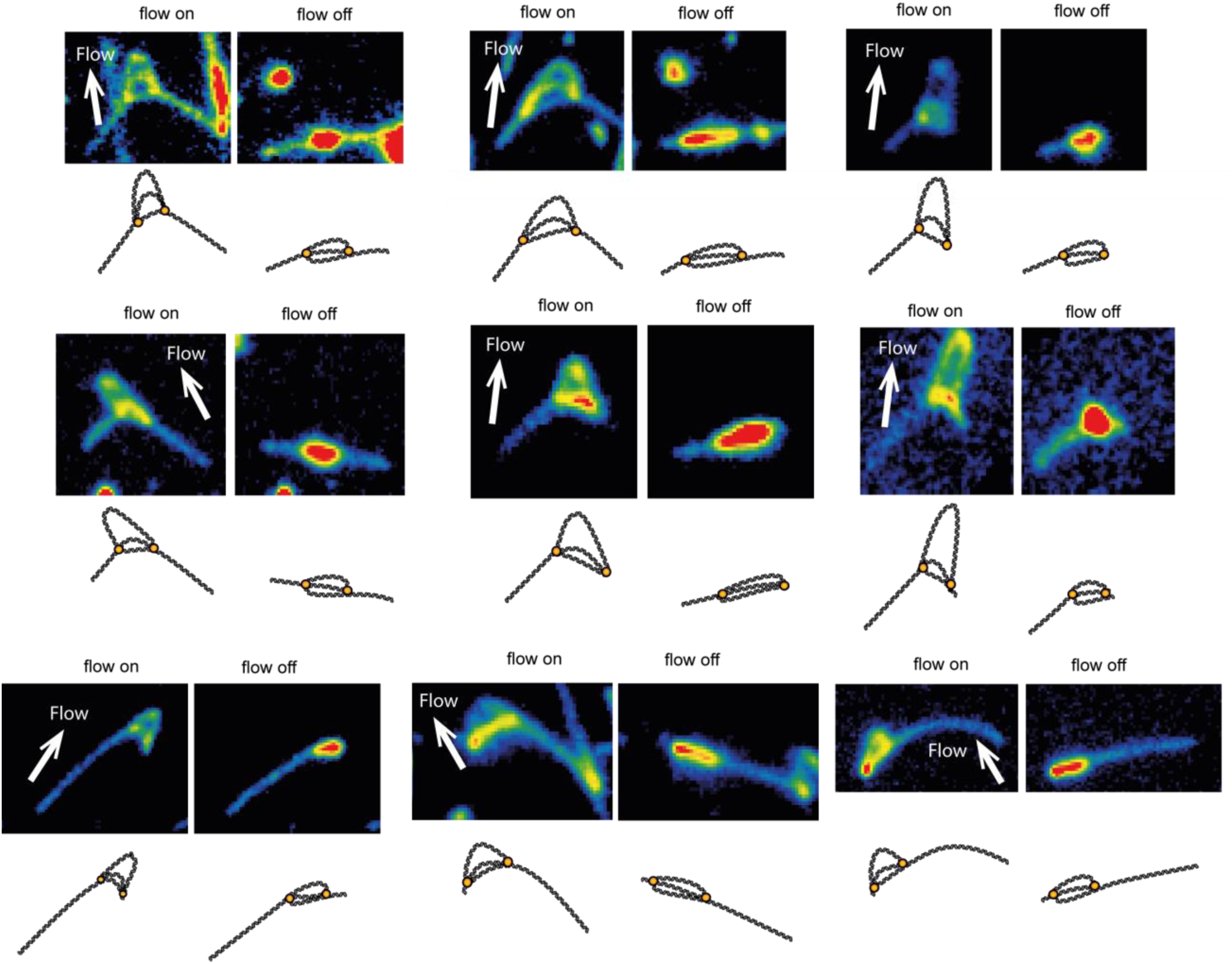
Additional examples of Z loops. Snapshots (top) and the corresponding schematics (bottom) of Z loops that were observed on different DNA molecules. By flow application (direction indicated by the white arrows), the three parallel dsDNA molecules that are linearly stretched along the DNA molecule are visualized.

**Extended Data Figure 3.**
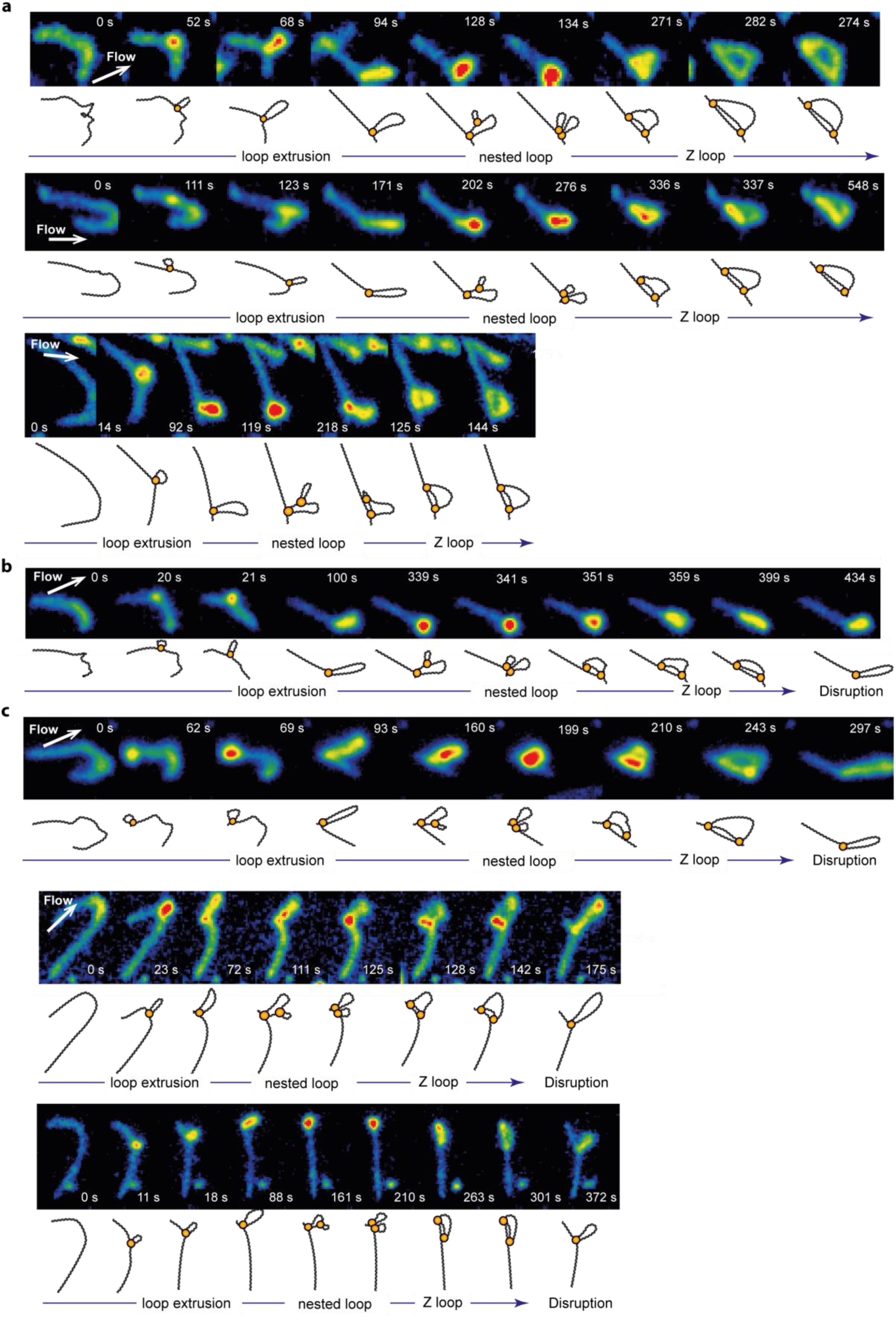
DNA visualizations during Z loop formation. **a, b, c,** Additional examples of series of snapshots (top) and the corresponding schematics (bottom) showing transitions from single-loop extrusion, to nested-loop formation, to Z-loop extension. Initially, a single loop is extruded and enlarged until it approached the tethered end of DNA. Subsequently, the single loop became further compacted, turning into the nested loop, followed by a gradual elongation into a Z loop. In the events of Z loop disruption **(b**,**c**), the structure reverted to a single loop, which was located either (**b**) at the position where the initial single loop had formed (See also Fig. 2B in main text) or (**c**) at the newly formed edge of the Z loop. The latter indicates that disruption occurred by release of the first condensin, after the second one had extended to the leading edge of the Z loop structure.

**Extended Data Figure 4.**
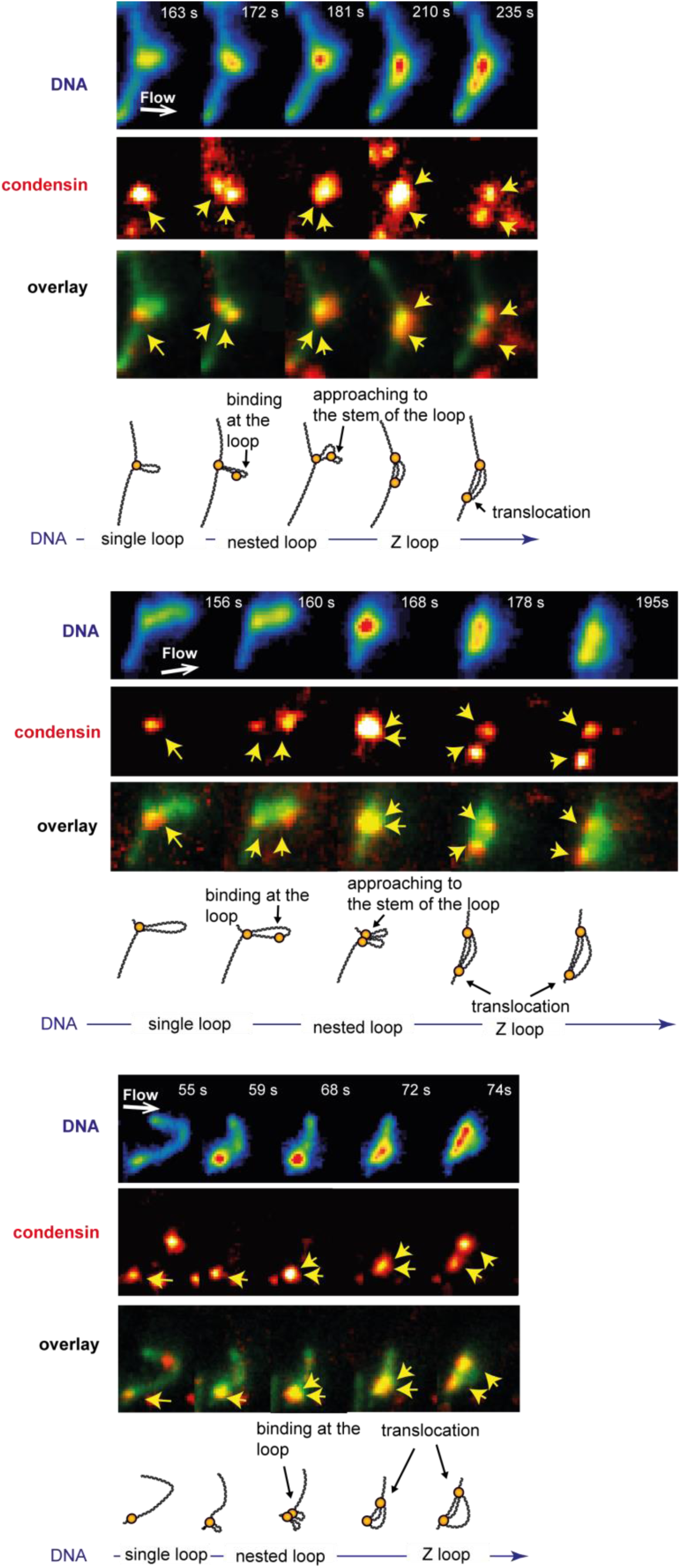
Co-visualization of DNA and condensins during Z loop formation. Additional examples of snapshots of SxO-stained DNA (top), ATTO647N-labeled condensin (middle), and their overlay (bottom) showing the locations of two condensins and the simultaneous transitions of DNA conformations during Z loop formation. A second condensin binds at the location within the loop that is created by the first condensin, and approaches to the first one while compacting the single loop further, whereupon one of the two condensins moves away from the other one as Z loop extends. Schematic diagrams underneath the images provide visual guidance. Arrows denote the location of the two condensins.

**Extended Data Figure 5.**
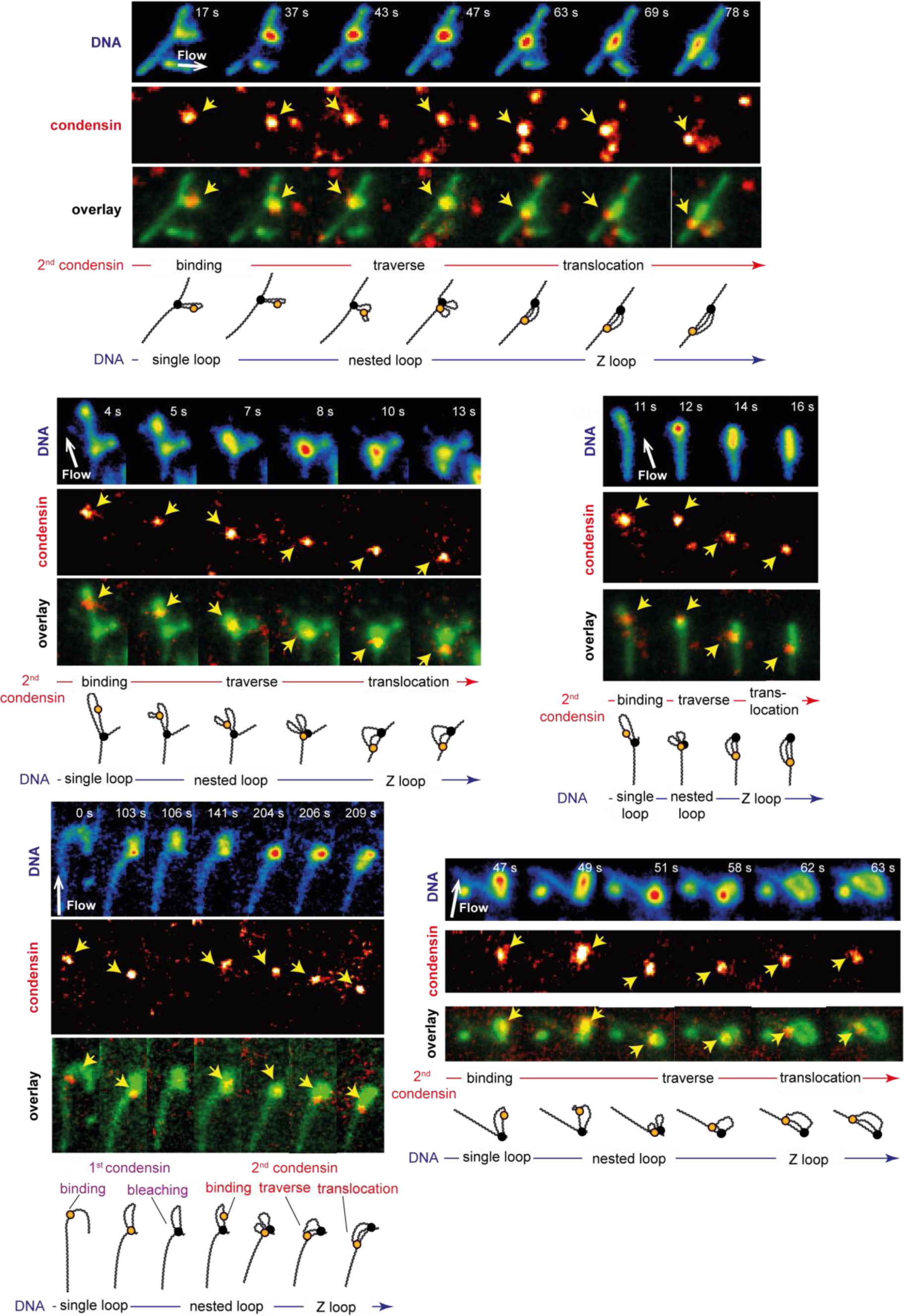
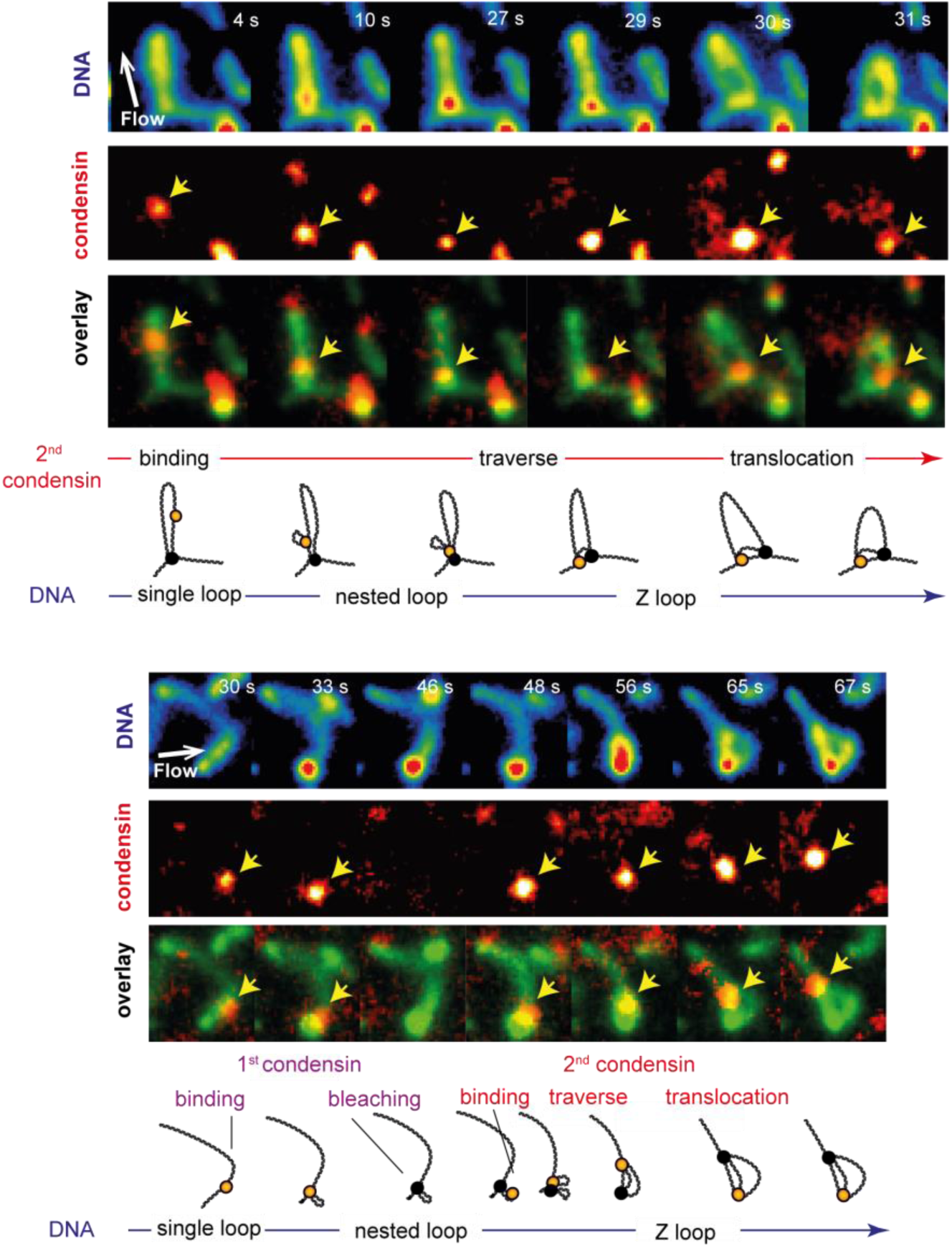
Co-visualization of DNA and condensins during Z loop formation, demonstrating condensin traversals. Additional examples of snapshots of SxO-stained DNA (top), ATTO647N-labeled condensin (middle), and their overlay (bottom) tracing the locations of the second condensin during Z loop formation. The fluorescence signal from the first condensin that extruded the single DNA loop was photobleached before the binding of the second condensin. Schematic diagrams underneath the images provide visual guidance.

**Extended Data Figure 6.**
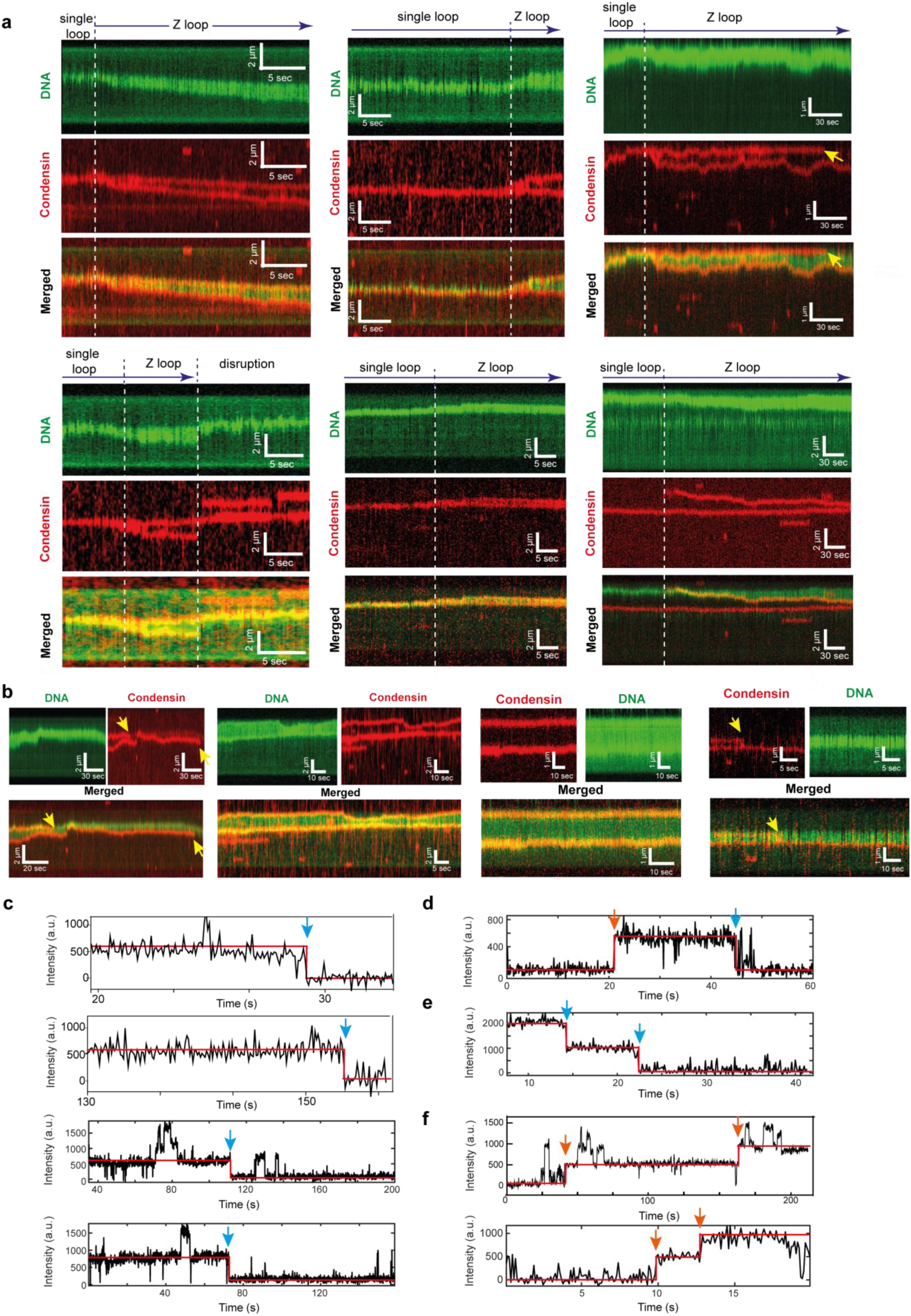
Kymographs and bleaching curves of condensins during Z loop formation on DNA. **a, b**, Additional examples of kymographs of SxO-stained DNA, ATTO647N-condensin, and their merge. The molecules in (**a**) shows a gradual extension of a single loop into a Z loop, and the simultaneous divergence of the locations of the two condensins that were previously co-localized near the single-loop region, into the edges of the Z-loop. Panel (**b**) shows condensins localized and co-fluctuating with the edges of Z-loop. Yellow arrows in (**a, b**) indicate the single-step photobleaching of the ATTO647N signals, while the Z loop remained stable. **c**, Additional examples of time traces of fluorescence intensities of individual ATTO647N-labeled condensin complexes near the edges of Z loops that bleached in a single step-wise manner. Note that occasionally an additional intensity is observed for short periods, which is due to the temporary binding of an additional condensin. **d**, Single-step binding and the consequent bleaching event of the same condensin for a DNA molecule for which the initial single loop was extruded by a non-labeled condensin, allowing us to monitor only the second condensin. **e**, Two-step bleaching events observed for a case where both condensins bleached while the Z loop of DNA remained. **f**, Two-step binding event observed upon initiation of Z loop formation. Blue arrows denote the bleaching events; orange arrows denote the arrival of the condensins.

**Extended Data Figure 7.**
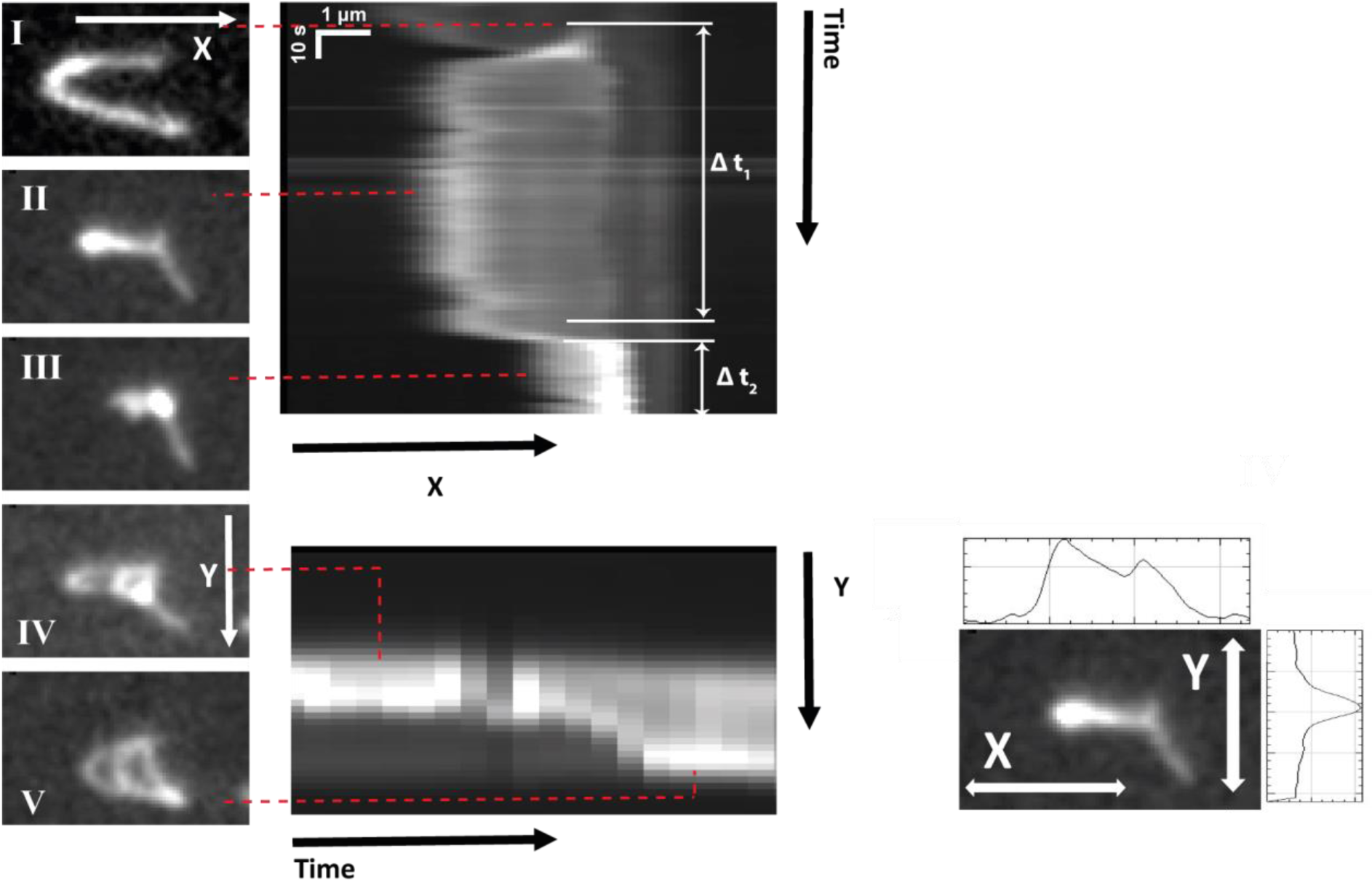
Expanded kymographs for the DNA molecule shown in Fig. 2C. Left panels: Snapshots of during Z-loop formation for the DNA molecule shown in Fig. 2C. Middle panels: Fluorescence intensity kymographs for this molecule along the axis parallel to the extruded single loop (X) and along the axis connecting both ends of the DNA tether (Y). These kymographs show a transition of a DNA conformation from a (I) bare DNA to (II) single loop, to (III) nested loop, to a (IV, V) gradual extension of a Z loop (cf. left panels). Such kymographs were used to estimate the time lags and the rate of nested loop plotted in Fig. 3F,G. Right panel: fluorescence intensity profiles extracted along the X and Y axis.

**Extended Data Figure 8.**
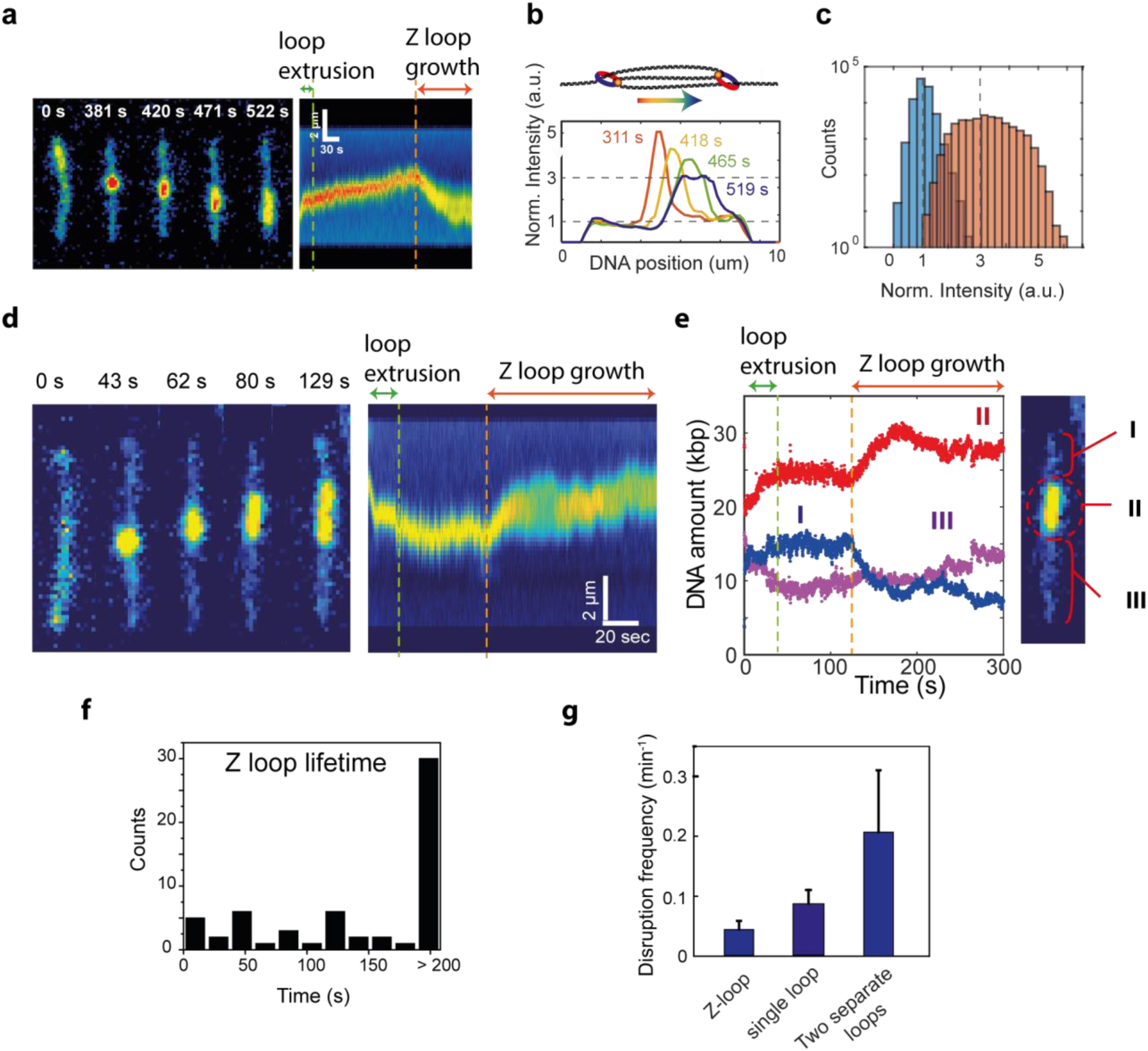
Quantitative characterization of Z loop formation in the absence of buffer flow. **a**, Snapshots and kymographs showing the formation of a single loop and the subsequent extension into a Z loop in the absence of flow. **b**, Time evolution (from orange to navy) of fluorescence intensity profiles extracted from five different lines of the kymograph in (A). Each plot is normalized to the average DNA intensity in the non-looped region. During Z loop formation, fluorescence intensity profiles initially displays a sharp peak, characteristic for a single loop, which progressively broadens, until it levels off at about 3 times of the intensity of the dsDNA outside the loop – consistent with three parallel dsDNA molecules. **c**, Normalized fluorescence intensity distributions extracted from fully stretched Z loops for the regions inside (orange) and outside (blue) the Z loops, showing an average value at 1-fold and 3-fold the single-DNA value, consistent with Z loops consisting of three dsDNA stretches in parallel. **d**, additional snapshots (left) and kymograph (right) showing the formation of a single loop and the subsequent extension into a Z loop, and **e**, the amounts of DNA extracted from the kymograph for the loop region (II) and the regions outside the loop (I and III). Dashed lines indicate the termination of the single loop growth (green), and the initiation of the Z-loop formation (orange). The rates of single loop extrusion and Z loop growth were extracted by a simple linear fit to the linear rise in the first 10 seconds of data in traces like shown here. **f**, Plot of the lifetime of extruded DNA loops (defined as the time between the start of Z loop extension and disruption into a single loop). About 50% of the extruded DNA loops did not release within the observation window of 200 seconds. **g**, Frequency of disruption events per DNA molecule that had, a Z loop, a single loop, or two separate loops – showing that Z loops are the most stable, even more so than single loops or two separate loops. Error bars are SD.

**Extended Data Figure 9.**
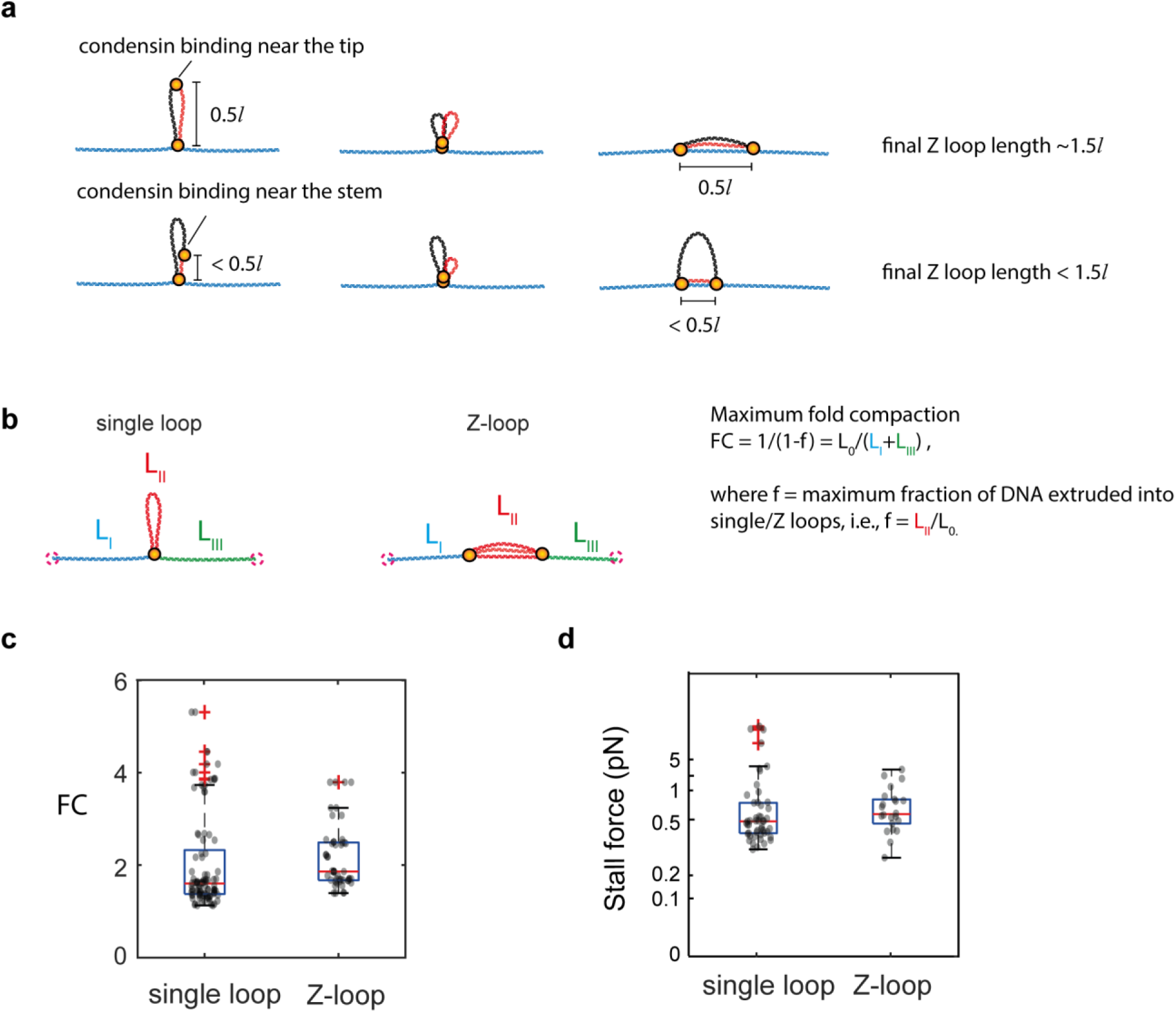
DNA compaction by Z loops versus single loops. **a**, Visual explanation on how the final length of Z loop relates to the initial single loop size *l*. **b**, Visual explanation on definition of maximum fold compaction (FC).^2^ **c**, Maximum fold compaction (FC) and **d**, the corresponding stall force (estimated from the known force-extension relation^3^) for single loops (N=49) and Z loops (N=22). The similar FC shows that two condensins forming a Z loop do not linearly compact DNA more than two condensins forming separate single loops.

**Extended Data Figure 10.**
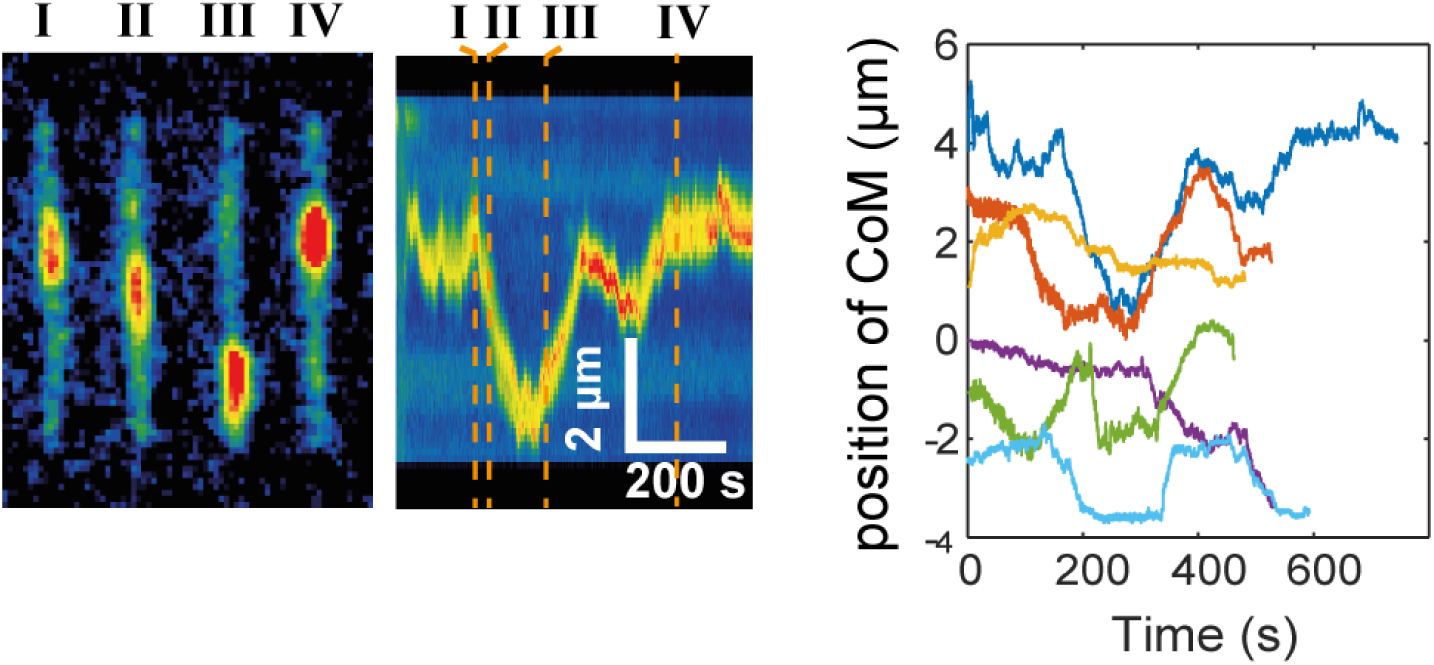
Z loops can diffuse along the DNA tether. Snapshots (left) and kymograph (middle) of a DNA molecule showing a Z loop that exhibits random diffusion along the DNA tether, presumably due to a tug of war between the two partly slipping condensin motors. The right panel shows time traces of center-of-mass positions of Z loops that diffuse along DNA (N=6).

## Supplementary Video captions

**Supplementary Video 1**. Movie showing interactions between two DNA loop that are located far apart. A newly forming DNA loop that was extruded by the second condensin grew in intensity (i.e. loop size) at the expense of the original one, which concomitantly shrank in size over time.

**Supplementary Video 2**. Movie showing a Z-loop structure that consists of three dsDNA stretches that are connected in parallel.

**Supplementary Video 3**. Movie showing the transition of DNA conformations from a bare DNA, to a single extruded loop, to a nested loop, to a Z loop. Note that in between of 186s and 398s (where frames were cut) DNA stayed in its conformation of a fully extruded single loop.

**Supplementary Video 4**. Movie with overlaid channels of fluorescence from SxO-stained DNA (green) and ATTO647N-labeled condensin (red), showing a single loop that is extruded by the first condensin, binding of the second condensin within the loop, merging of two condensins accompanied by the formation of nested loop, followed by diverging of two condensins as the Z loop expands.

**Supplementary Video 5**. Movie with overlaid channels of fluorescence from SxO-stained DNA (green) and ATTO647N-labeled condensin (red), showing the single loop that is extruded by the first condensin, that was bleached before the binding of the second condensin at the DNA-loop region, whereupon the second condensin approached the DNA-loop stem, followed by a linear translocation along the DNA outside of the initial loop.

**Supplementary Video 6.** Movie with overlaid channels of fluorescence from SxO-stained DNA (green) and ATTO647N-labeled condensin (red), showing the binding of the second condensin at the DNA-loop region, and its subsequent approach to the DNA-loop stem, where the first (photobleached) condensin resides, followed by the linear translocation along the DNA outside of the initial loop. Note that in this rare uncommon case, the tip of the single loop got stuck on the surface for a moment, allowing us to better localize the movement of the second condensin.

**Supplementary Video 7**. Movie with overlaid channels of fluorescence from SxO-stained DNA (green) and ATTO647N-labeled condensin (red), showing condensin localization to the Z-loop edges and their spatial fluctuations that correlates with the edges of the Z loop upon buffer flow application to the left and right.

**Supplementary Video 8 and 9.** Movies showing two-side-pulling Z-loop where DNA is reeled in symmetrically from both sides, leading to a simultaneous expansion of the Z loop to both directions.

**Supplementary Video 10**. Movie showing a Z-loop that fills the unextruded DNA gap between two single loops.

